# Memory consolidation and improvement by synaptic tagging and capture in recurrent neural networks

**DOI:** 10.1101/2020.05.08.084053

**Authors:** Jannik Luboeinski, Christian Tetzlaff

## Abstract

The synaptic-tagging-and-capture (STC) hypothesis formulates that at each synapse the concurrence of a tag with protein synthesis yields the maintenance of changes induced by synaptic plasticity. This hypothesis provides a biological principle underlying the synaptic consolidation of memories that is not verified for recurrent neural circuits. We developed a theoretical model integrating the mechanisms underlying the STC hypothesis with calcium-based synaptic plasticity in a recurrent spiking neural network. In the model, calcium-based synaptic plasticity yields the formation of strongly interconnected cell assemblies encoding memories, followed by consolidation through the STC mechanisms. Furthermore, we find that the STC mechanisms have an up to now undiscovered effect on memories – with the passage of time they modify the storage of memories, such that after several hours memory recall is significantly improved. This kind of memory enhancement can provide a new principle for storing information in biological and artificial neural circuits.

## Introduction

In biological neural systems, memories have a wide repertoire of dynamics; most importantly, they can be encoded, stored, recalled, and consolidated. While these dynamics are relatively well-explored at the behavioral and brain-region level [22, 24, 31], the underlying synaptic and neuronal processes remain mainly elusive.

Most generally, learning describes the ability of humans and other animals to obtain knowledge about an entity. This knowledge or information is stored as a memory. The encoding of such a memory in a neural network is commonly assumed to happen in the way as described by Hebb in his seminal work [25, 35, 48]: A group of recurrently connected neurons that receives the information by an external input starts to fire stronger than the rest of the network. This increased firing yields strengthening of the efficacy of the recurrent synapses within this particular group such that a so-called Hebbian cell assembly is formed, which represents a memory of the input. On the other hand, low firing rates typically cause weakening of connections between neurons, which can lead to either disruption or refinement of a cell assembly. Strengthening and weakening of synapses at timescales relevant to memory is the result of long-term synaptic plasticity [2, 6, 9].

Long-term synaptic plasticity creates long-lasting changes of the synaptic efficacy. To become strengthened, synapses undergo a cascade of molecular processes that leads to an increase in the number of postsynaptic receptors, which is called long-term potentiation (LTP, [2, 5, 12, 75]). Analogously, for weakening, another cascade of processes yields a decrease in the number of receptors, which is called long-term depression (LTD, [2, 5, 75]). The signaling cascades of both LTP and LTD are triggered by the calcium concentration in the postsynaptic spine. The spiking activities of the pre- and the postsynaptic neurons drive the calcium concentration and, by this, determine whether long-term potentiation or long-term depression of the synaptic efficacy is induced [6, 33, 34, 43, 72]. In general, long-term synaptic plasticity consists of at least two different phases. Changes of the synaptic efficacy in the early phase last for several hours, while efficacy changes in the late phase can be maintained for several days [1, 12]. The transfer from the early to the late phase has been described by the synaptic-tagging-and-capture (STC) hypothesis [28, 65]. Following the STC hypothesis, the transfer depends on the stimulation of the specific synapse as well as on the stimulation of other synapses at the same postsynaptic neuron. More precisely, the transfer at a synapse occurs if the synapse is tagged, which means that it is primed for transfer, and if proteins necessary for the late phase are abundant or have been synthesized. The tagged synapse then “captures” proteins causing the transfer to the late-phase by increasing for instance the number of receptor slots at the postsynaptic site [65]. The formation of a tag at a synapse is related to its own early-phase change, while protein synthesis depends on the early-phase state of many synapses [17, 28, 43, 65]. The total or effective synaptic efficacy of a synapse consists of the sum of the early- and late-phase contribution [17, 43].

In general, consolidation of memories means the progressive transfer of memories into a state in which they stay stable over long time intervals [22, 24, 54]. There a two major categories of memory consolidation: systems consolidation and synaptic consolidation [22]. The basic idea of systems consolidation is that a memory is transiently stored in the hippocampus and possibly transferred to the neocortex, in which it is maintained for a longer period. The question, whether a memory is first encoded in the hippocampus and then transferred to the neocortex or whether the encoding of a memory occurs simultaneously in both regions (multiple trace theory, [22, 56]), is subject to an ongoing debate. In both cases, however, the newly formed memory has to be initially consolidated such that systems consolidation can set in. This initial consolidation process, named synaptic consolidation, is related to local molecular and cellular processes at individual neurons and synapses [16, 23, 49].

The STC hypothesis provides a potential explanation of the neuronal and synaptic processes underlying the synaptic consolidation of memories [22, 28, 65], which is supported by several theoretical studies focusing on single synapses or feed-forward networks [8, 17, 43, 75, 83]. However, a clear link between the STC hypothesis and memory consolidation is still missing because, as discussed above, the encoding of memories in neural circuits is mainly associated with strongly recurrently connected groups of neurons (cell assemblies).

In this study, we developed a theoretical model of recurrently connected spiking neurons with the synaptic efficacies being altered by calcium-dependent synaptic plasticity and the core mechanisms of the STC hypothesis. The individual components of the implemented model reproduce various plasticity phenomena as the ones described above [10, 28, 42, 57, 58, 66, 69, 73, 74, 80, 82] and for verification we matched the temporal evolution of individual synapses in our model with experimental data. Our simulations show the synaptic and neuronal dynamics underlying the formation of a memory representation, its recall, and its long-term development. The latter indicates that the STC mechanisms in a recurrent circuit yield the consolidation of a memory. Finally, the simulations and analytical results suggest a new implication of the STC mechanisms on memory representations, which is the enhancement of the storage of the memory. This enhancement exhibits a new type of memory dynamics, which could be beneficial for biological as well as for artificial memory systems.

## Results

We aimed to set up a biophysically plausible network model of the neuronal and synaptic dynamics underlying synaptic consolidation. For this, based on previous studies [17, 33, 43], we utilized a synaptic model that integrates the local calcium dynamics that trigger synaptic plasticity and the mechanisms of synaptic tagging and capture (Fig. 1a). We used this synaptic model in conjunction with a leaky integrate-and-fire model, axonal delay, and a finite synaptic time constant to cover the most important aspects of signal generation and transmission in neural networks with parameters based on experimental findings (see Methods, [30, 77]). Before introducing the details of the network model, first, we will present the key principles of the synapse model and compare its dynamics to experimental data.

**Figure 1:**
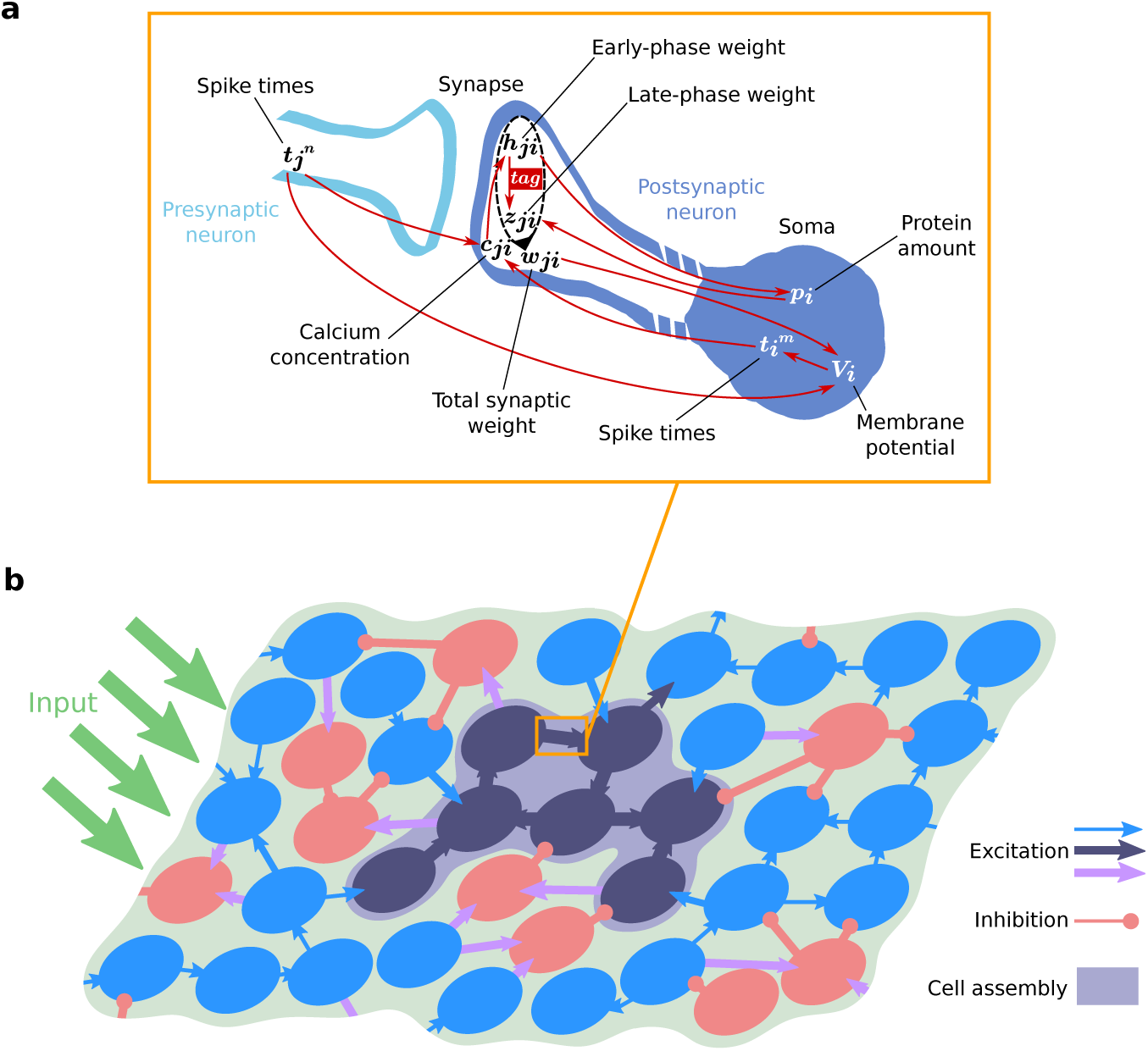
Schematic of the synaptic and network model. **(a)** The synaptic model integrates the interplay between various mechanisms of calcium-dependent synaptic plasticity and the STC hypothesis. For more details see the main text. **(b)** Schematic of a part of the neural network that consists of excitatory (blue and dark blue circles) and inhibitory neurons (red circles) and receives external input from other brain areas (green arrows). Only synapses between excitatory neurons (blue and dark blue arrows) undergo plastic changes by the processes shown in (a). A Hebbian cell assembly represents a memory and consists of a group of strongly interconnected neurons (dark blue).

### Comparison of synapse model with experimental data

In general, the model assumes that the membrane potential of a neuron determines its spiking dynamics that drives together with presynaptic spiking the postsynaptic calcium concentration (Fig. 1a). The calcium concentration determines the occurrence of early-phase LTP and early-phase LTD, represented by changes in the early-phase component or weight of the synaptic efficacy. Large changes of the early phase weight trigger the formation of a synapse-specific tag. A sufficient body of early-phase changes at many synapses of the postsynaptic neuron triggers protein synthesis. Once an adequate amount of proteins is available and the synapse is tagged, the late phase component or weight of the synaptic efficacy is altered; thus, the synapse “captures” proteins. The sum of the early- and late-phase weight yields the total synaptic efficacy that determines the magnitude of postsynaptic potentials arriving at the neuron, influencing its membrane potential. The interplay between these different processes is investigated in standard plasticity induction experiments [65, 69, 70]. In these experiments a strong tetanus stimulation (STET) is used to induce late-phase potentiation, while weak tetanus stimulation (WTET) is used for early-phase potentiation only. For late-phase depression, a strong low-frequency stimulus (SLFS) is used, while for early-phase depression, a weak low-frequency stimulus (WLFS) suffices. As proof of concept of our synaptic model, we reproduced the outcome of these experiments by considering a single plastic synapse and applying similar stimulation protocols (see Supplementary Fig. S1 for details). The resulting time traces of the synaptic efficacy in response to the four induction protocols (Fig. 2) match the findings from experiments as discussed above and from previous theoretical studies [8, 17, 43, 83]. These results indicate that our synaptic model provides a reasonable description of the underlying biomolecular dynamics. Thus, in the next step, we will introduce our analysis of the synaptic and neuronal dynamics in a recurrent neural network, which are based on the synaptic model.

**Figure 2:**
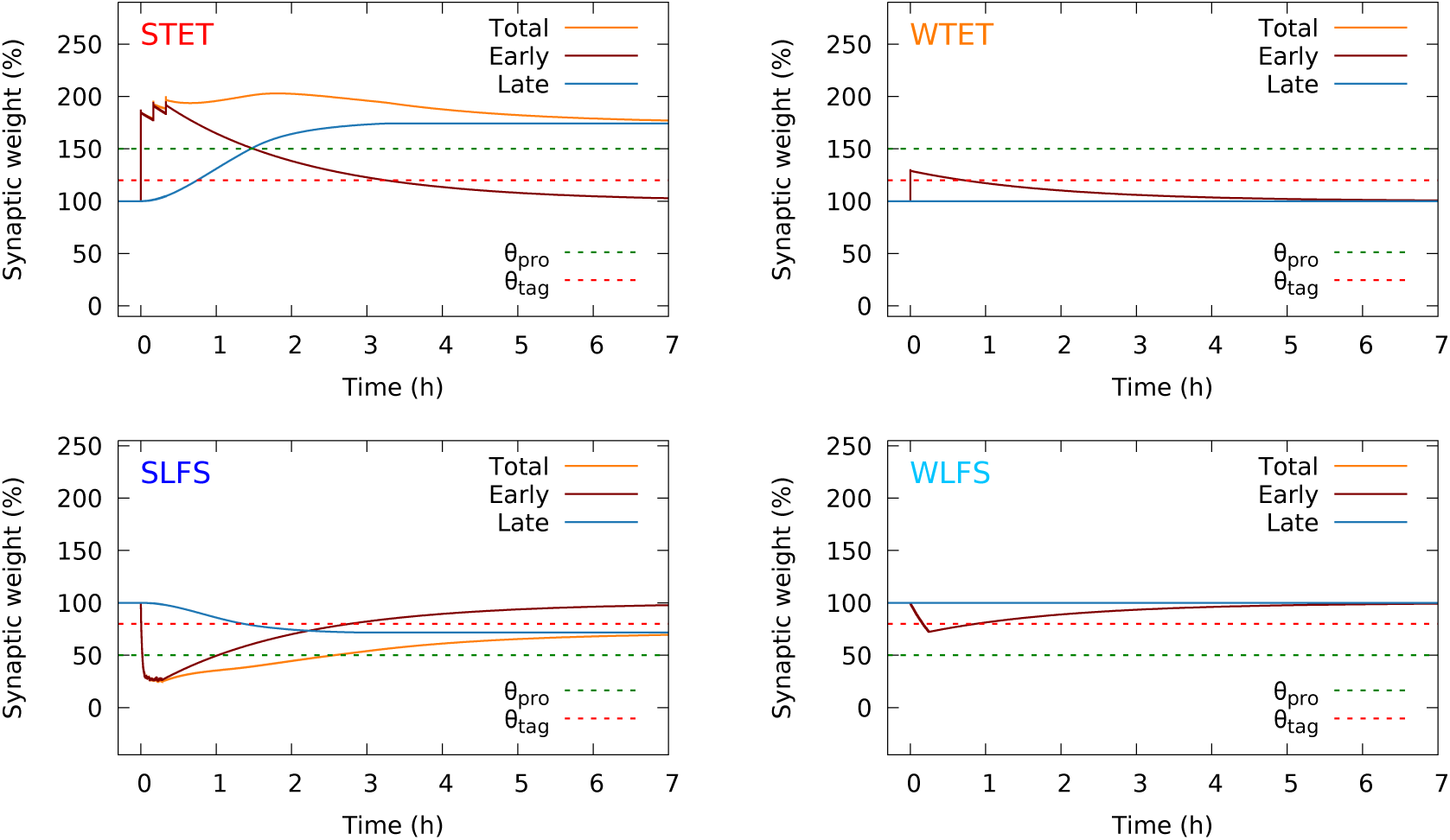
Induction of late-phase potentiation (STET), early-phase potentiation (WTET), late-phase depression (SLFS), and early-phase depression (WLFS) at a single synapse with standard protocols. The late-phase weight (blue line) is only changed by strong stimulation (STET, SLFS). The early-phase weight (red line) is affected by all protocols. WTET and WLFS suffice to drive the early-phase weight across the threshold of tag formation (*θ*_tag_, dashed red line) but not across the threshold of triggering protein synthesis (*θ*_pro_, dashed green line). The total weight or synaptic efficacy (orange) is the sum of early- and late-phase weight. For graphical reasons, the late-phase weight has been shifted (see Methods).

### Network model with synaptic consolidation enables functional memory representations

Employing our synaptic model, we simulated a patch of hippocampus consisting of 2000 neurons with 20% being inhibitory and a 10% average probability of two neurons being connected by a synapse (Fig. 1b). Synapses between excitatory neurons (blue, dark blue) feature plasticity mechanisms as described above. Inhibitory neurons provide feedback inhibition; their connections are non-plastic (purple, red). All neurons in the patch received additional inputs from outside the network, which have particular characteristics for the encoding and recalling of a memory. During learning, a specific stimulus is provided to a group of neurons and should trigger the formation of particularly strong connections between these neurons, which then represent a Hebbian cell assembly or memory (Fig. 1b, dark blue). During recall, only a subset of the group of neurons that received the learning stimulus received specific external stimulation.

To investigate the synaptic and neuronal dynamics implied by the STC mechanisms, throughout this study, we focused on two time spans. We evaluated the state of the neural network around 10 seconds after the learning stimulus to analyze the short-term dynamics and after around 8 hours to investigate the long-term effects of the STC mechanisms.

The learning stimulus consisted of three subsequent pulses of 0.1 seconds each, strongly activating a random subset of neurons. As expected, the stimulus caused tremendous changes of the synaptic efficacy of diverse synapses in the network (compare Fig. 3a with b). Synapses associated with the stimulated neurons (first 150 neurons Fig. 3a-c) mainly experienced LTP (red). The synapses between stimulated neurons (black box in Fig. 3a-c) are strengthened particularly, indicating the correct formation of a cell assembly. By contrast, synapses between non-stimulated neurons underwent LTP as well as LTD (blue). After 8 hours, the synaptic changes between non-stimulated neurons fade, such that mainly changes associated with the stimulated neurons and, thus, with the cell assembly remain (Fig. 3c). To validate that the formed cell assembly encodes a memory, next, we tested the ability of recall. For this, we applied a recall stimulus that activates for 0.1 seconds 50% of the neurons that were stimulated by the learning signal, and analyzed the resulting spread of activation within the network (Fig. 3d, e). The externally stimulated neurons (‘as’) have the highest activity. By the strong recurrent connections in the cell assembly, the average activity of the non-stimulated neurons in the cell assembly (‘ans’) is significantly higher than the activity of the remaining non-stimulated neurons (‘ctrl’). This is not the case for control stimulation applied before learning. In other words, the activity of the recall stimulus spreads via the stimulated neurons to the other cell assembly neurons and yields a pattern completion or recall. Please note that, under basal (standby) conditions, the mean firing rate of all neurons remains in the typical physiological range for the hippocampus of 0.5–1.0 Hz ([50]; see also Supplementary Fig. S3). These results indicate that calcium-dependent synaptic plasticity and the STC mechanisms together enable the encoding of memory and its maintenance for several hours in a recurrent neural network. Thereby, our findings support the idea that the STC mechanisms account for the synaptic consolidation of memories.

**Figure 3:**
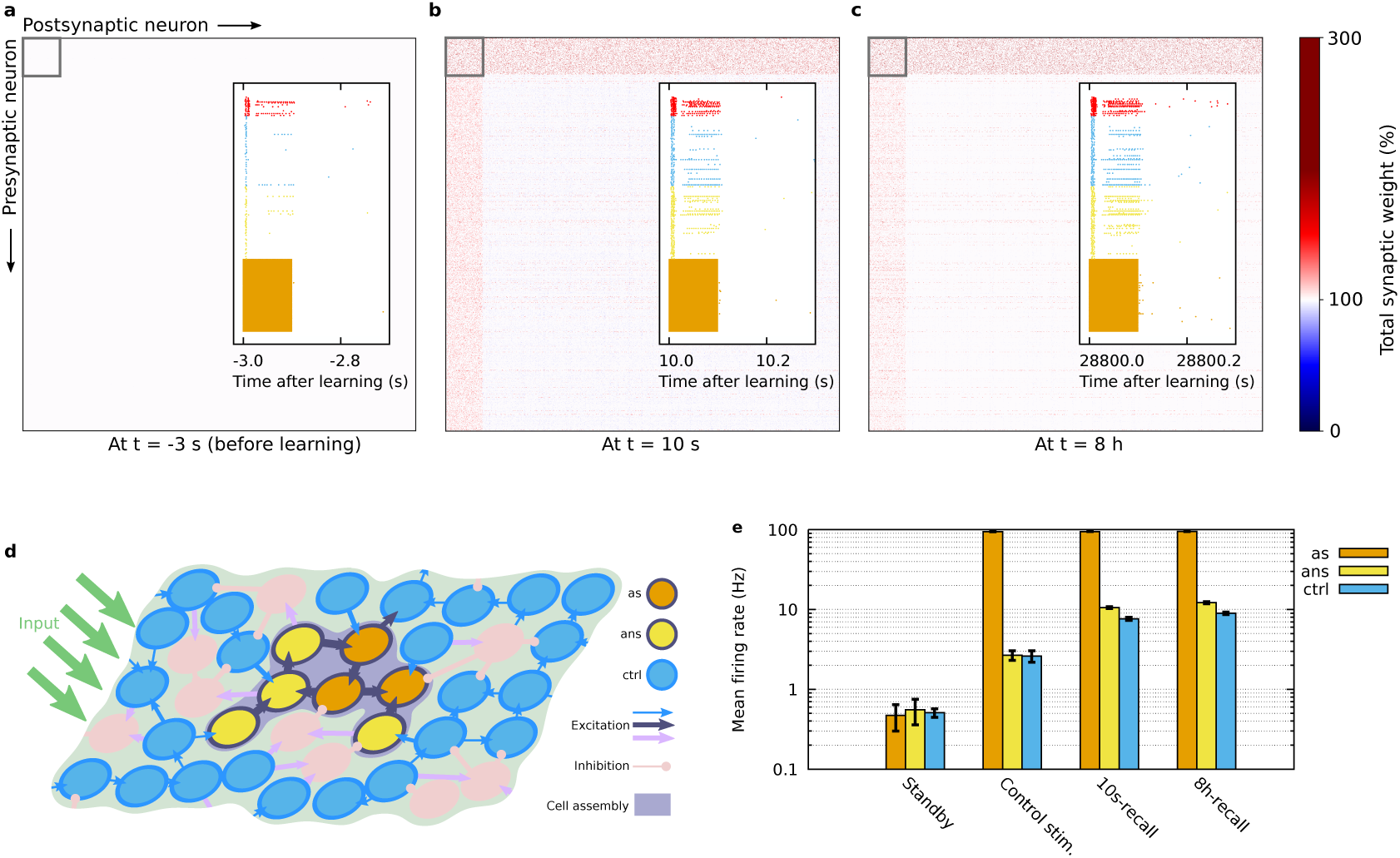
Formation and temporal development of a Hebbian cell assembly. Panels (a–c) show the matrix of total (early- plus late-phase) synaptic weights sorted by the stimulated neurons (first 150): **(a)** before learning, **(b)** around 10 seconds after learning, and **(c)** around 8 hours after learning. The insets show the spiking during recall stimulation of the neurons in the cell assembly (colors as in (d,e)) and of a random 5% fraction of the neurons in the remaining excitatory (blue) and inhibitory (red) populations. **(d)** Schematic of the three subpopulations during recall stimulation: the fraction of externally stimulated cell assembly neurons (‘as’), the fraction of cell assembly neurons that are not stimulated (‘ans’), and the remaining excitatory neurons acting as control group (‘ctrl’). **(e)** The mean firing rates of the three subpopulations defined in (d), before learning and without recall (standby), before learning during control recall stimulation, upon recall 10 s after learning, and upon recall 8 h after learning. Error bars indicate the standard deviation across the respective subpopulation. Synaptic weights and mean firing rates were averaged over 10 trials. Parameters: *w*_ie_*/h*_0_ = 4, *w*_ii_*/h*_0_ = 4.

### Memory functionality depends on network inhibition and size of the representation

In the following, we investigate the influence of different parameters such as inhibition and cell assembly size on the learning, recalling, and consolidation of memory representations.

In general, one role of inhibition in neural networks is to prevent the network activity from runaway or epileptiform dynamics. This kind of dynamics is characterized by neurons constantly exciting each other and firing at the maximum possible rate. On the other hand, if inhibition is excessively strong, neuronal activity is confined, resulting in a more or less silent network. We adjusted the level of inhibition in our network by varying the coupling strength from inhibitory neurons to excitatory neurons (*w*_*ie*_) and the coupling strength from inhibitory to inhibitory neurons (*w*_*ii*_). For every set of these parameter values, we simulated the network dynamics during learning and recall as described before. At two different points in time (10 s and 8 h after learning), we evaluated the memory functionality that consists of learning, consolidation and recall of a memory by measuring the recall quality. To measure the recall quality, we used two different quantities (cf. Methods): on the one hand, the pattern completion coefficient *Q* describes by how much the activity in the non-stimulated part of the cell assembly is raised during recall as compared to the activity of the control neurons; on the other hand, the mutual information *MI*_*v*_ describes how similar the network activity state during recall is to the network activity state during learning. We quantitatively defined memory functionality by an average pattern completion coefficient of *Q* ≥ 0.03. This criterion demands that, upon stimulation of half of the assembly neurons, the other half of the assembly be activated much stronger than the background neurons, which remain at a low activity level (cf. Fig. 3e).

High values of the I→I coupling strength together with low values of the I→E coupling strength imply a low level of inhibition, which impedes in our network model the functionality of a memory (see Fig. 4a,c for recall performance 10 s after learning and Fig. 4b,d for 8 h after learning). We expect that this is due to the runaway dynamics discussed before. If a network is in such an epileptiform state, a recall stimulus cannot exclusively activate the neurons that belong to a cell assembly, as also all the control neurons become active overshadowing memory recall. On the other hand, low values of the I→I coupling strength together with high values of the I→E coupling strength lead to extensive levels of inhibition suppressing the neuronal activity. This also impedes memory functionality, because memory recall becomes impossible, as any spread of activity within the network will immediately be suppressed. As a result, the level of inhibition has to be within a specific regime to enable learning, consolidation and successful recall of memories (indicated by the red box; *Q* ≥ 0.03). Within this regime, the level of inhibition seems to be sufficiently high to prevent the overly activation of control neurons, while it remains low enough to allow the desired activation of the non-stimulated neurons in the cell assembly. Please note that this regime could be related to the network state of loosely-balanced excitation and inhibition [20]. The regime of memory functionality is the same either 10 s or 8 h after providing the learning stimulus. However, the higher values of *Q* and *MI*_*v*_ after 8 h compared to the 10 s case indicate a positive influence of long-term dynamics on the memory functionality. We further investigate and discuss this result in the next section.

**Figure 4:**
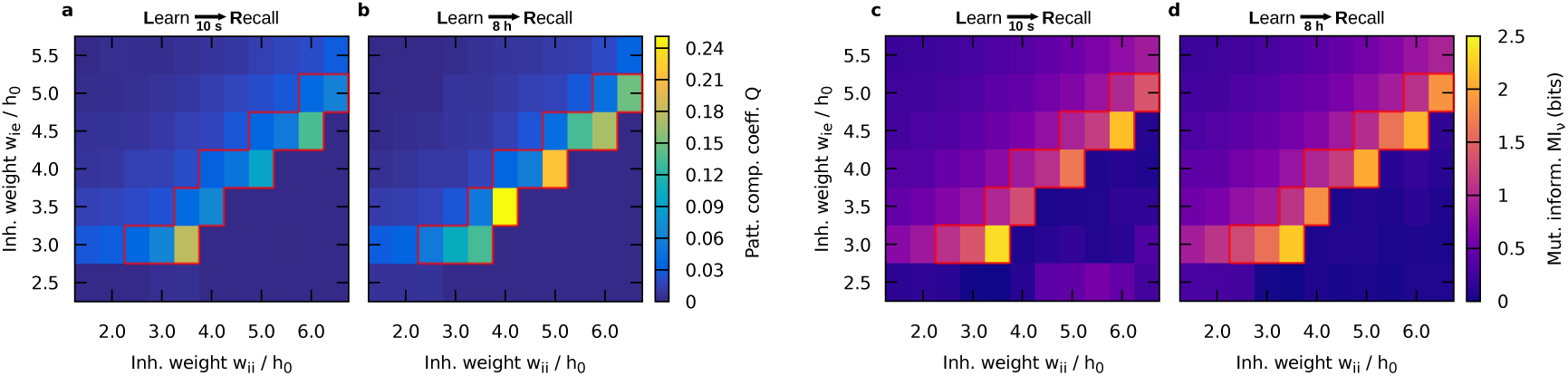
Quality of the recall 10 s and 8 h after learning as a function of the I→E (*w*_*ie*_) and I→I (*w*_*ii*_) coupling strengths. Sufficient memory functionality is found within a certain regime, marked by the red box (*Q* ≥ 0.03). Two measures of recall quality are shown: **(a**,**b)** the pattern completion coefficient *Q* (with non-significant values set to zero, see Methods); **(c**,**d)** the mutual information *MI*_*v*_ between the neuronal activities in the network during learning and during recall. Parameters: *n*_CA_ = 150.

As further parameter, we examined the influence of the size of the cell assembly on the memory functionality (Fig. 5a,b). We controlled the size by varying the number of neurons being the subgroup that is stimulated by the learning stimulus. Following our definition of memory functionality, requiring that the coefficient *Q* be greater or equal than 0.03, we found that learning and recall is only possible if the cell assembly is large enough. In the following, we will focus on a particular set of parameter values for the inhibition (*w*_ie_*/h*_0_ = 4, *w*_ii_*/h*_0_ = 4). All derived conclusions apply to all sets within the specific regime of inhibition discussed before (red box in Fig. 4). For small cell assemblies (here about 150 neurons), the group of neurons is too small to exhibit functional pattern completion (threshold indicated by the dotted red line in Fig. 5a,b). For large cell assemblies (here above 500 neurons), the activity of the cell assembly becomes self-sustained. This means that after learning the neurons within the cell assembly stay active, preventing the typical dynamics of memories we are looking for. Moreover, the measures of the recall quality exhibit local maxima that could emerge due to “resonance effects” in the interaction between the excitatory and inhibitory population of the network, which depend on the number of stimulated excitatory neurons. The occurrence of local maxima cannot be explained by the relationship between the mean firing rate and the cell assembly size, because this relationship is strictly monotonic and does not feature any maximum (Supplementary Fig. S3). Finally, similar to Fig. 4, the pattern completion coefficient and mutual information become higher 8 h after learning compared to 10 s after learning for a large range of cell assembly sizes, which will be further examined in the next section.

**Figure 5:**
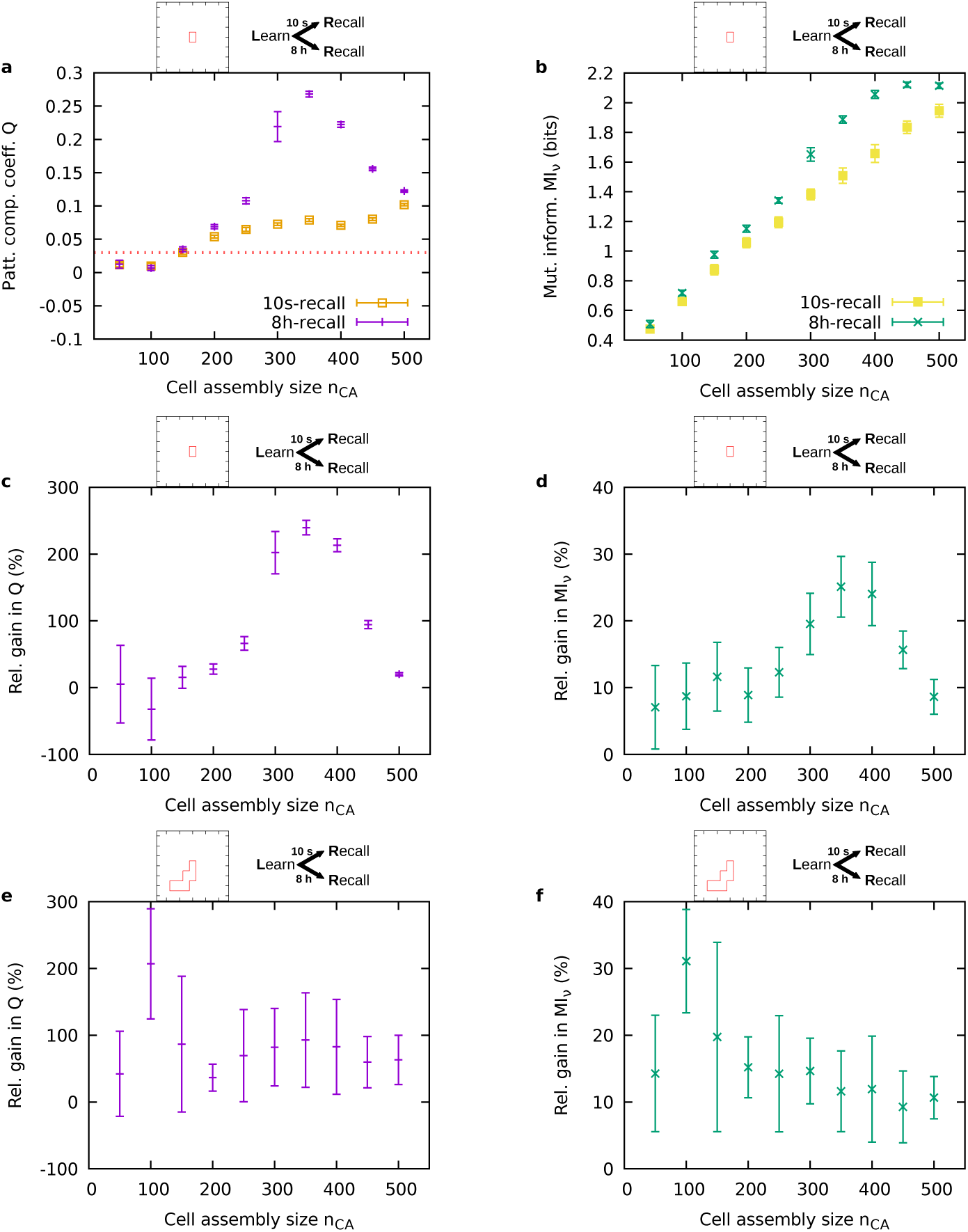
Recall quality 10 s and 8 h after learning and resulting relative gain as a function of the cell assembly size. **(a)** Pattern completion coefficient 10 s (orange) and 8 h (purple) after learning. The dotted red line indicates the threshold for memory functionality (*Q* ≥ 0.03). **(b)** Mutual information of the neuronal activities 10 s (yellow) and 8 h (green) after learning. **(c)** Relative gain in pattern completion coefficient between 10 s and 8 h after learning. **(d)** Relative gain in mutual information of the neuronal activities between 10 s and 8 h after learning. (a-d) Parameter values: *w*_ie_*/h*_0_ = 4, *w*_ii_*/h*_0_ = 4. **(e)** Average relative gain in pattern completion coefficient across multiple parameter sets yielding memory functionality. **(f)** Average relative gain in mutual information of the neuronal activities across multiple parameter sets yielding memory functionality. (e,f) All parameter sets *w*_ie_*/h*_0_ ∈ [2.5, 5.5], *w*_ii_*/h*_0_ ∈ [1.5, 4.0] with *Q*(10s-recall) ≥ 0.03. Error bars indicate the standard deviation across ten trials.

### Consolidation improves memory recall

Comparing the recall quality of the cell assembly 10 s after learning with the recall quality 8 h after learning for different paradigms (Figs. 4 and 5a,b), we found that 8 h after learning the system generally features an improved recall quality. In other words, the specific state of the cell assembly after eight hours, resulting from the interplay between calcium-dependent synaptic plasticity and the STC mechanisms, seems to facilitate recall. To elucidate the magnitude of the improvement effect, we computed the relative gain in recall quality between the state 10 s after learning and the state 8 h after learning and present results for the particular inhibition setting that we chose before (5c,d). It becomes obvious that improvement in recall quality can occur by more than 200% with respect to the pattern completion coefficient and by as much as 30% with respect to the mutual information.

To test the robustness of the improvement effect across the specific inhibition regime identified before, we averaged *MI*_*v*_ and *Q* over the range of parameter settings of this regime and calculated the relative gain in recall quality (Fig. 5e,f). For the chosen inhibition setting *w*_ie_*/h*_0_ = 4, *w*_ii_*/h*_0_ = 4 (Fig. 5c,d) as well as for the averaged results (Fig. 5e,f), we observed a strictly positive gain across all cell assembly sizes (one data point was slightly negative but non-significant). Positive gain means that 8 h after learning the performance was better than 10 s after learning. Thus, for a wide range of parameter settings, the passage of time yields an improvement of the recall quality.

What are the underlying principles that lead to this improvement of the recall quality? We could identify two processes, both being related to the STC mechanisms: an active and a passive improvement.

The *active improvement* is related to the dynamics of the early-phase weights. The recall stimulus provided 10 s after learning leads to only minor variations of the average early-phase weight in the cell assembly (red dashed line at ‘R’ in Fig. 6a). By contrast, 8 h after learning the recall stimulus triggers a significant increase of the average early-phase weight, which in turn results in a stronger total synaptic weight (red and orange line at ‘R’ in Fig. 6b). Thus, 8 h after learning the average total synaptic weight of the cell assembly increases during the recall that improves pattern completion or rather the recall performance (Fig. 5). By comparing the state of the neural network between both time points, we find that the average early-phase weight before the 10s-recall resides on a much higher level than before the 8h-recall (Fig. 6c). This difference in early-phase weight before recall stimulation could explain the different dynamics of the early-phase weights during the recall. Considering the mathematical formulation of the model, one can see that the change of the early-phase weight depends on the distance of the actual early-phase weight *h* from its initial or equilibrium value *h*_0_. Hence, larger changes occur if the early-phase weight is closer to the equilibrium state, while the changes remain smaller if the early-phase weight is close to its maximum value. Thus, the learning stimulus “pushes” the early-phase synaptic weight into a regime in which the subsequent recall cannot trigger further strengthening. However, with the passage of time the early-phase weight decays (while the late-phase weight increases) until it reaches the vicinity of its initial value (Fig. 6b,c). In this regime a recall stimulus can again trigger an increase of the early-phase weight, supporting pattern completion. The detailed weight distributions for all cases presented in Fig. 6 are shown in Supplementary Fig. S4. To further scrutinize the relation between the early-phase dynamics during recall and the improved recall quality, we performed simulations in which we switched off or blocked early-phase plasticity after learning. If we compare the resulting recall performances with the simulations with early-phase plasticity (Fig 7), for both measures *Q* and *MI*_*v*_, we do not find an influence of the blockage during 10s-recall but during 8h-recall. These results further support our finding that the dynamics of the early-phase weights within the cell assembly are one essential part underlying the improvement of the recall quality.

**Figure 6:**
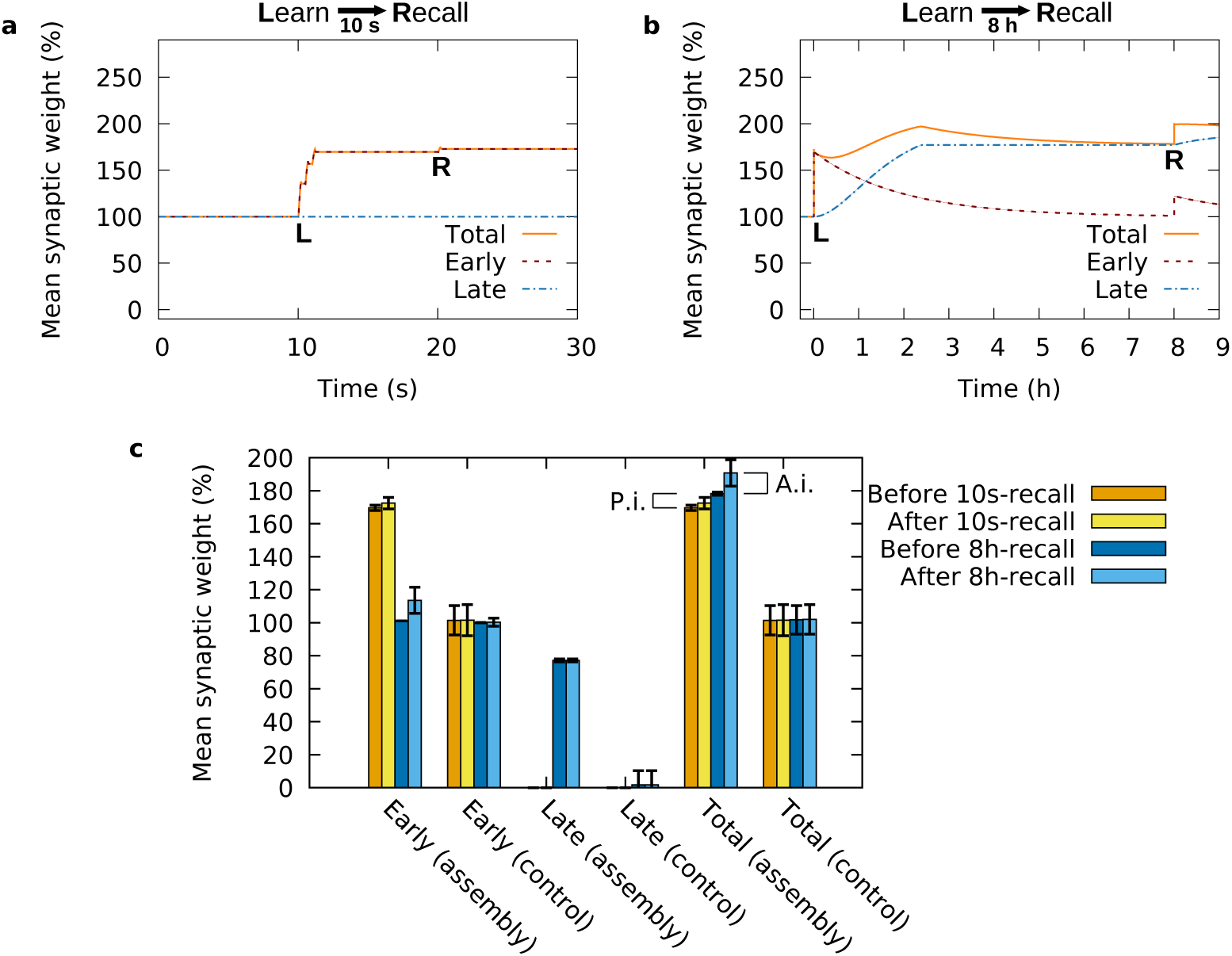
Synaptic dynamics related to memory formation, consolidation and improvement. **(a)** During the first seconds after the learning stimulus (‘L’), the average early-phase weight (red) of the synapses connecting the stimulated neurons increases, while the average late-phase weight remains constant (blue). A recall stimulus (‘R’), provided 10 s after the learning stimulus, does not have a significant effect on the synaptic weights. **(b)** Several hours after learning, the average early-phase weight decays while the late-phase increases such that the average total synaptic weight remains on a high level. Providing a recall stimulus 8 h after learning triggers an increase of the average early-phase weight and, thus, of the total synaptic weight. **(c)** Mean early- and late-phase weights within the assembly and within the non-assembly (control) population immediately before and immediately after the 10s- and 8h-recall. A.i.: active improvement component (difference between dark and light blue); P.i.: passive improvement component (difference between orange and dark blue). Error bars show the standard deviation across the subpopulation. In (a,b), the errors are too small to be visible. Parameter setting: *w*_ie_*/h*_0_ = 4, *w*_ii_*/h*_0_ = 4, *n*_CA_ = 150.

**Figure 7:**
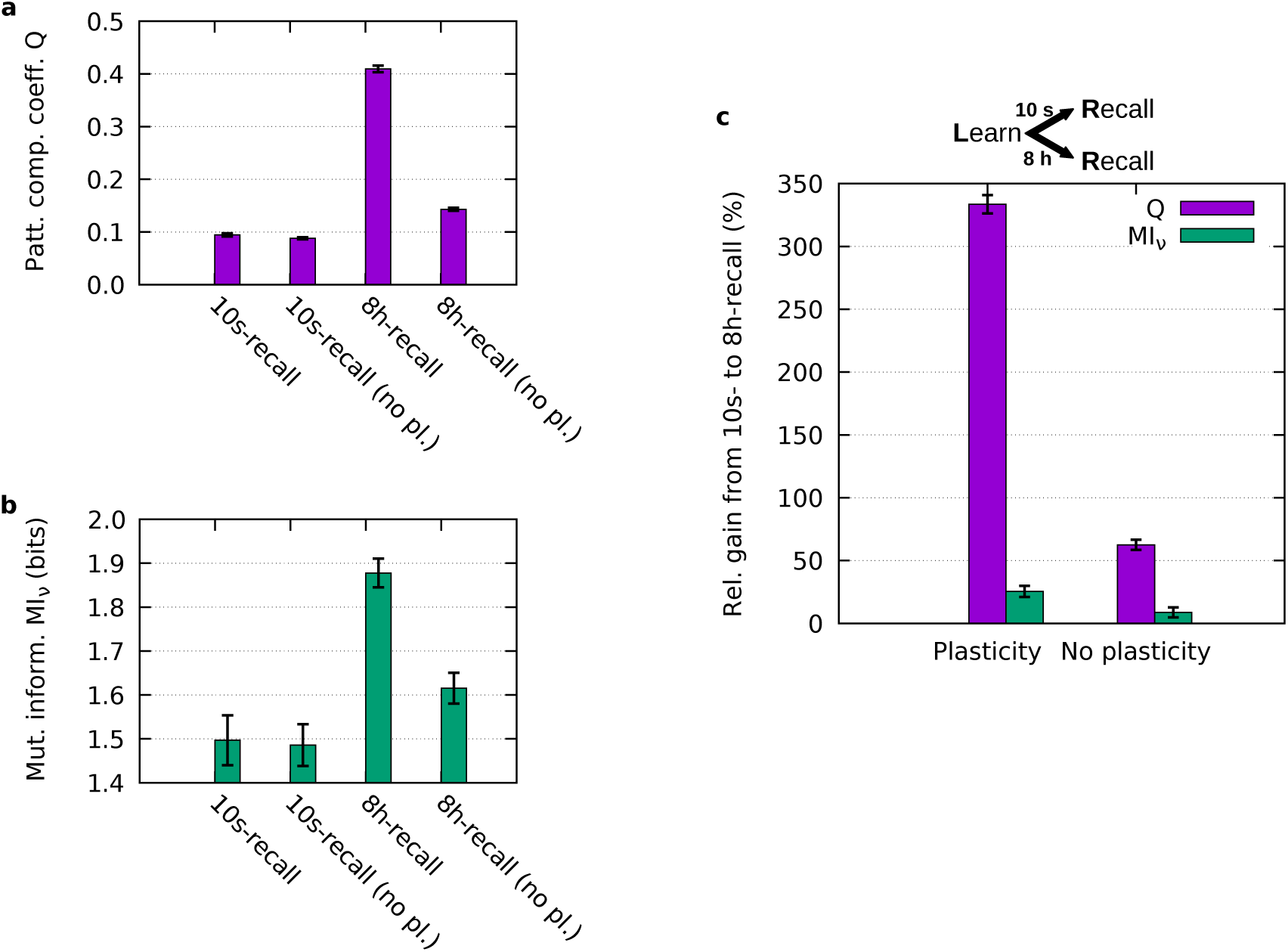
Recall quality measured for recall 10 s and 8 h after learning with and without early-phase synaptic plasticity during recall. The effect of early-phase synaptic plasticity on the total improvement is significant (compare 8h-recall with 8h-recall no pl.; active improvement), but it does not cover the complete level of improvement (compare 10s-recall with 8h-recall no pl.; passive improvement). **(a)** Pattern completion coefficient *Q*; **(b)** Mutual information *MI*_*v*_. **(c)** Relative gain in *Q* and *MI*_*v*_ from 10s-recall to 8h-recall. Average across 10 trials. Error bars indicate standard deviation across trials in (a,b) and the propagated error in (c). Parameter setting: *w*_ie_*/h*_0_ = 4, *w*_ii_*/h*_0_ = 4, *n*_CA_ = 350 (value of the maximum in Fig. 5c,d).

The other part of the improvement of the recall quality is related to the dynamics of the late-phase weights within the cell assembly. We refer to this part as *passive improvement*. As expected, 8 h after learning the STC mechanisms yield a decrease of the average early-phase weight (red line in Fig. 6b) accompanied by an increase of the average late-phase synaptic weight in the assembly (blue line) that indicates the process of synaptic consolidation. However, if we compare the average total synaptic weight within the assembly before the 10s-recall with the one before the 8h-recall (Fig. 6c), we identify an increase providing a second component underlying the improvement of recall quality also indicated by the relative gain in *Q* and *MI*_*v*_ without early-phase plasticity (Fig. 7). Hence, although the dynamics of the early-phase weights (active improvement) explains the gain in recall quality to a large extent, there is a remaining part that is related to the dynamics of the late-phase weights (passive improvement).

### Parameter dependency of the passive improvement

We have found that the mean total weight of a cell assembly can increase over time, and refer to this phenomenon as passive improvement (cf. Fig. 6c). This effect is elicited by the mechanisms of synaptic tagging capture and occurs if the mean late-phase weight 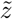 after eight hours becomes higher than the mean early-phase weight 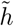 after learning. To investigate and predict the circumstances under which this effect is elicited, namely, different settings of late-phase-related constants, we considered an analytical approach that we present in the following.

To investigate the parameter-dependency of consolidation and improvement of a cell assembly, we analytically solve the linear first-order differential equation describing the dynamics of late-phase LTP (cf. Eq. 14) with a variable coefficient introduced by the time-dependent amount of proteins (Eq. 15). Since L-LTP is vastly predominant between cell assembly neurons, we neglect here the L-LTD dynamics.

Thus, the analytical expression for the mean late-phase weight after time *t* after learning in the case of persistent tags and protein synthesis is:

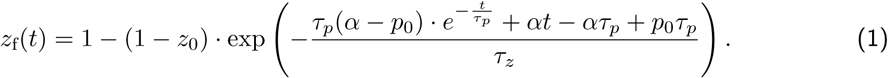

depending on the protein time constant *τ*_*p*_, the late-phase time constant *τ*_*z*_, the initial protein amount *p*_0_, the initial late-phase weight *z*_0_, and the protein synthesis rate *α*. This solution can be further simplified under the condition that the initial late-phase weight *z*_0_ and the initial protein amount *p*_0_ are zero. The mean increase in late-phase weight due to the STC mechanisms is then:

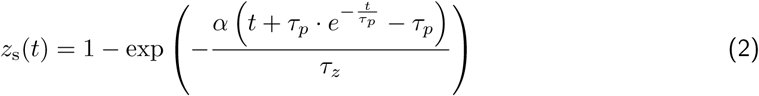

For Eq. 2, we considered that the synapses within the cell assembly are always tagged, triggering changes of the late-phase weights. However, after learning the early-phase weight decays (Fig. 6b) and falls at a certain point in time below the tag-sufficient threshold. Thus, first, we have to calculate the decay of the mean early-phase weight and, then, use it to determine when the tag vanishes. The average decaying early-phase weight follows

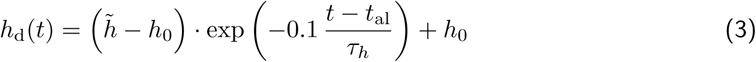

depending on the mean early-phase weight after learning 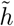, the time after learning *t*_al_, the initial weight *h*_0_, and the early-phase time constant *τ*_*h*_. The time *t*_al_ is the time after learning (for instance, 10 seconds) at which the mean learning-induced increase in early-phase weight, 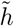, is measured.

Setting *h*_d_(*t*) to the value of the tagging threshold *θ*_tag_ + *h*_0_ yields the point in time *t*_tag_ when the synaptic tags vanish on average:

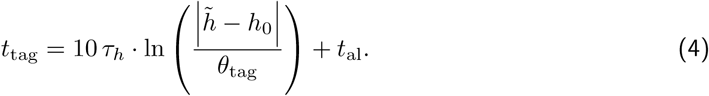

The synaptic tags typically vanish before the synthesis of new proteins stops and, thus, *t*_tag_ determines the end of late-phase changes, such that the average late-phase weight is described by

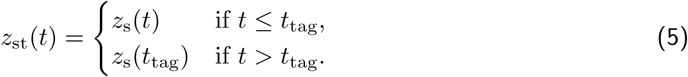

Hence, for parameter sets yielding *t*_tag_ *<* 8 h (cf. Eq. 4), the mean late-phase weight before the 8h-recall is the same as by the time the tags vanished (see also Fig. 6b, blue line):

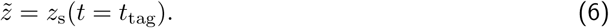

In Fig. 8a we plot the mean weights within the cell assembly, computing the late phase via the analytical function in Eq. 5, using our numerical results for early-phase LTP during learning and recall, and computing the decay of the early-phase weight as detailed in Eq. 3. We find that this analytical approach predicts very well the mean weight dynamics of our numerical simulations (dashed gray line in Fig. 8a; Fig. 6b).

**Figure 8:**
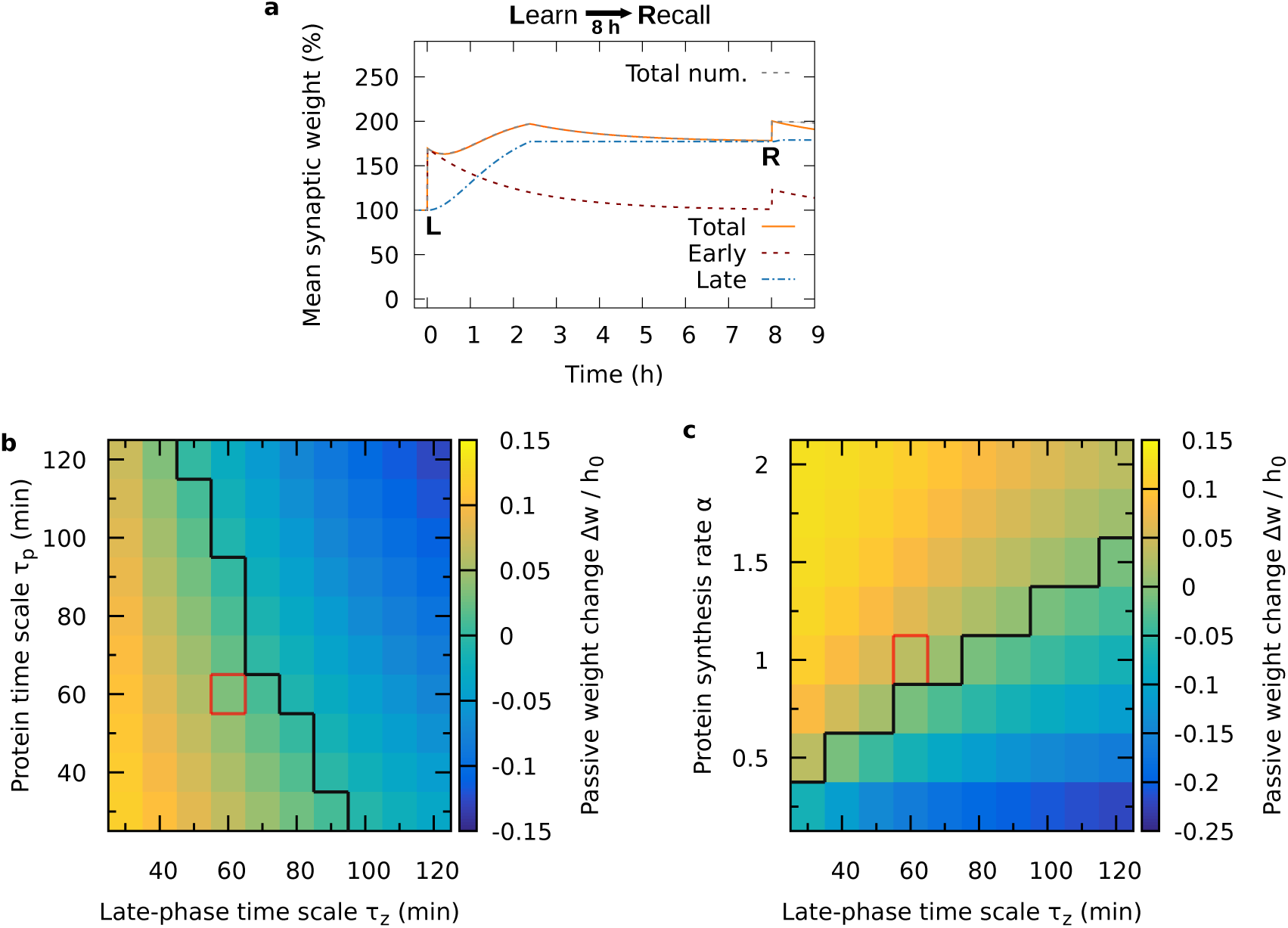
Analytical calculation of the long-term mean weight dynamics. **(a)** The early-phase values immediately after learning (‘L’) and after recall (‘R’) were taken from numerical simulations and inserted into the analytical equations for the late-phase dynamics (Eq. 5) and the early-phase decay (Eq. 3). The curves are in very good match with the simulated results, the total weight of which is shown here for comparison by the dashed gray line (cf. Fig. 6b). **(b**,**c)** Varying the time constants for late-phase dynamics *τ*_*z*_ and protein amount *τ*_*p*_ (b), as well as the protein synthesis rate *α* (c), shows that for slower time scales and lower synthesis rate, the passive weight change becomes negative, i.e., there is a passive deterioration instead of improvement. The black separatrix line demarcates the two regimes of improvement and deterioration. The red box indicates the parameter values used in our numerical studies.

When we compare the mean total weight 8 h after learning *w*(*t* = 8 h) with the mean total weight 10 s after learning *w*(*t* = 10 s), we obtain an expression for the passive weight change realized by the STC mechanisms, thus, of the passive improvement:

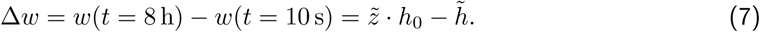

If Δ*w* > 0, the system shows a passive improvement with the passage of time, which is given for a wide range of parameter values. For instance, by considering different values of the protein synthesis parameters, namely the time constant of the protein amount *τ*_*p*_ (Fig. 8b) and protein synthesis rate *α* (Fig. 8c), depending on the time scale of the late-phase dynamics *τ*_*z*_, we can determine the influence of protein synthesis on the improvement during recall. We find that, if the protein time scale becomes too high or the synthesis rate becomes too low, the passive weight change switches from improvement to deterioration. However, the protein dynamics can be much slower than in our numerical simulations (red box) and still the STC mechanisms yield an improvement. For the protein synthesis rate, there is also a wide range of values that gives rise to improvement. Thus, the encountered improvement effect is robust to parameter variations.

Please note that our analytical results lead to some predictions: if the time scale of the protein dynamics is manipulated, the resulting memory behavior should switch between improvement and deterioration. In addition, manipulating the speed of signaling cascades that are involved in late-phase LTP should have a similar effect. Taking this into account, our predictions can be useful to narrow down the set of possible proteins and processes involved in synaptic tagging and capture.

### Intermediate recall further amplifies memory improvement

There are several psychological studies that investigate hypermnesia/improvement by considering the memory strength after multiple recalls [27, 61, 79]. Therefore, we also investigated the impact of an intermediate recall on the memory dynamics. For this, we computed the gain in the pattern completion coefficient *Q* and in the mutual information *MI*_*v*_ between 10s-recall and 8h-recall after applying an additional, intermediate recall stimulus.

We varied the time of occurrence of this intermediate stimulus (Fig. 9). The gain in recall quality for early intermediate recall (minutes after learning) shows little difference to the gain without intermediate recall (data point at time zero). For late intermediate recall, the gain reaches values that are much higher compared to the case without intermediate recall. Hence, additional recall improves the recall quality even further, which is consistent with experimental findings [27, 61, 79].

**Figure 9:**
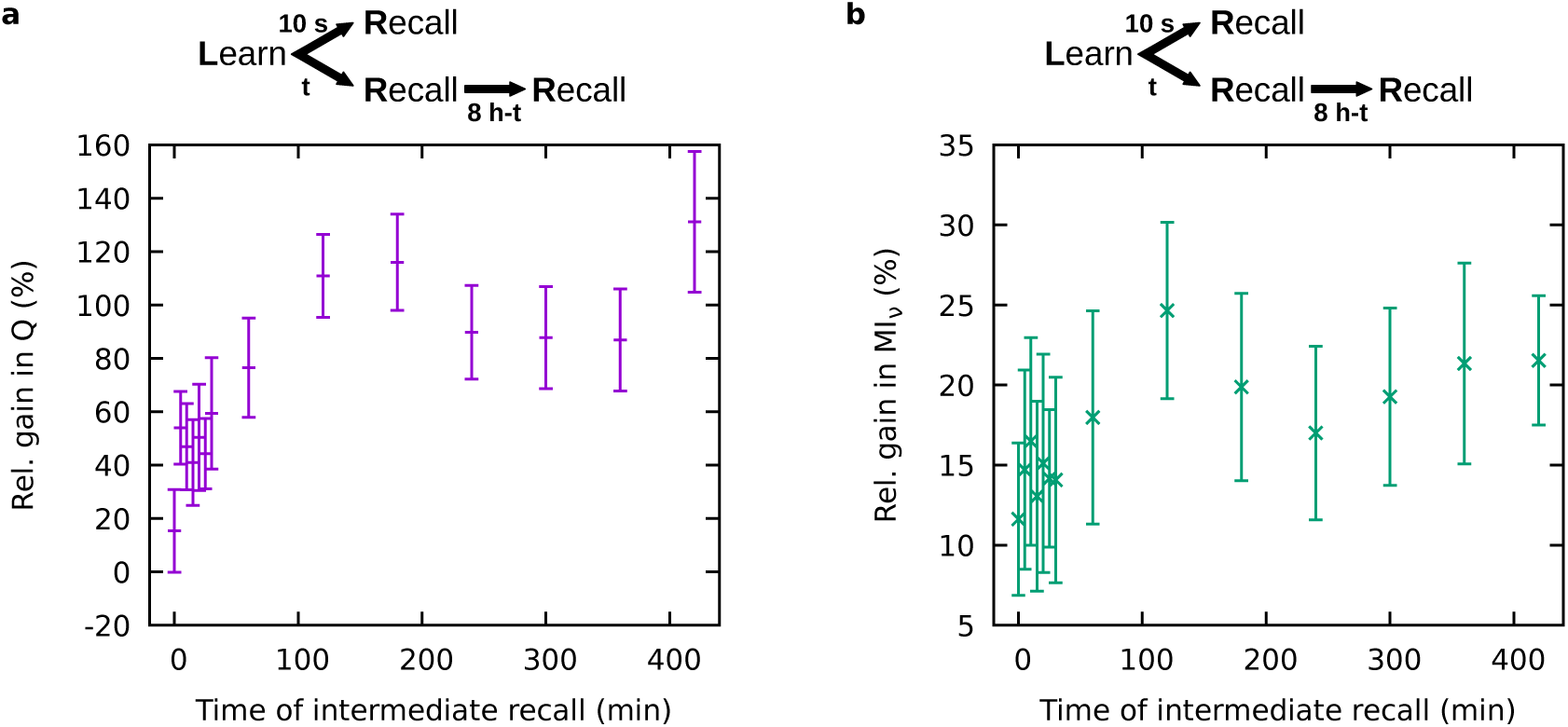
Relative gain in recall quality between 10 s and 8 h after learning, affected by an intermediate recall stimulus at varying times. **(a)** Gain in pattern completion coefficient *Q*; **(b)** gain in mutual information *MI*_*v*_. Parameter values: *w*_ie_*/h*_0_ = 4, *w*_ii_*/h*_0_ = 4, *n*_CA_ = 150. The data points at time zero show the case without intermediate recall (as in Fig. 5c,d). Error bars indicate the standard deviation across trials.

### Testing our predictions in experiments

Our model predicts an active and a passive improvement of memories by the synaptic tagging and capture mechanisms for a wide range of parameter values. In this section, we present some predictions that can be verified experimentally including experiments on the behavioral level, on the network level, and on the synaptic level.

At the behavioral or psychological level, our findings can be tested with subjects that need to learn a set of items, like words or pictures, and then recall the items after a very short time (around ten seconds). The performance of this recall should then be compared with the performance of the same subjects in a similar task but with other items, and recall after eight hours. Ideally, to be comparable to our results, there should not be any attempt to recall during the eight hours, which could be hard to manage for the subjects. However, at least trying to avoid intermediate recall would provide some new insights as compared to previous psychological studies that were based on intermediate recall [27, 61, 79].

In in-vitro networks, multielectrode arrays (MEAs) can be used to stimulate many neurons at the same time, as well as to measure their activity [46]. For our purposes, an MEA could be used to trigger plasticity, forming a strongly interconnected cell assembly. After the formation, a partial recall stimulus should be applied and the resulting activity be measured. The activity after 10s-recall (ten seconds after learning) and after 8h-recall (eight hours after learning) can then be compared as it was shown in Fig. 3e. If the activity in the cell assembly following 8h-recall is higher, then there will be experimental evidence for memory improvement. Another promising approach to test memory improvement at the network level would be to use optogenetic stimulation in combination with activity measurements through calcium imaging, which is an established method being very suitable for in-vivo experiments [4, 76], and can be tuned to deliver precise learning and recall stimulation [15, 68].

Our study also yields predictions on the synaptic level. To test this, first, synaptic potentiation has to be induced applying a strong ‘learning’ stimulus to a nerve fiber. Later application of a shorter ‘recall’ stimulus to the fiber should then cause a response that depends on the time that has passed since learning. Following our predictions, the response, which can be measured as a relative field excitatory postsynaptic potential (often abbreviated %fEPSP), should be larger eight hours after learning than ten seconds after learning, corresponding to the mean total synaptic weight shown in Fig. 6. Furthermore, after the application of two ‘recall’ stimuli, the response should even be higher, as we showed in the previous section.

Finally, in any experiment, blocking the dynamics that lead to early-phase plasticity should diminish the improvement effect significantly, because we predict that active improvement is mainly triggered by the early-phase dynamics (cf. Fig. 7c). Such experiments could be realized by using NMDA receptor antagonists or by blocking exocytosis before the presentation of a recall stimulus. In addition, blocking or slowing down late-phase-related signaling cascades and protein dynamics should, beyond a certain level, prevent passive improvement and thereby also diminish the total improvement effect (cf. Fig. 8b,c).

## Discussion

In this study, we showed with a recurrent network model that the mechanisms of synaptic tagging and capture [17, 28, 43, 65] can provide a biological basis for encoding, consolidation, and recall of memories. We found that these mechanisms also cause an improvement in the recall quality of memories, which is robust across a large space of parameter settings. Previous theoretical studies of STC focused on molecular pathways [75], single synapses [8, 17, 43, 83], and feed-forward networks [83], and thereby provided fundamental insights into the processes underlying the synaptic consolidation of memories. However, to the best of our knowledge, there have not yet been studies targeting the effects of STC in recurrent networks, which are essential for the encoding of memories. We set out to close this gap and we were able to characterize emergent effects that arise from the conjunction of synaptic tagging and capture with strongly recurrently connected groups of neurons (cell assemblies).

Detailed models describing the molecular dynamics underlying synaptic plasticity and STC have been shown to be compatible to and offer a complementary view on simpler statistical models [8, 17, 32, 33, 75]. We use a model for synaptic plasticity based on calcium dynamics because this approach captures a wide range of experimental findings, including spike-timing dependent phenomena and rate-coding behavior [10, 33, 42, 57, 58, 72, 73, 80, 82]. For STC, we use a model based on generalized plasticity-related proteins that account for the crucial aspects of tagging, capture and cross-tagging [17, 43, 65, 66, 69, 74]. Thereby, our model contains the essential features of a biologically more detailed model, while still being computationally tractable.

By varying the size of the cell assembly, we found different recall dynamics of the system. We found that there is a lower bound to the size, below which the amplification of recall stimulation does not suffice for functional recall. On the other hand, large cell assemblies show attractor behavior, meaning that after the application of a recall stimulus they become fully activated and stay activated. Investigating this attractor regime is beyond this study. Nevertheless, considering attractor dynamics in our model could reveal interesting implications, since long-term memory representations exhibiting attractor dynamics can serve as working memory [60, 63], possibly in conjunction with additional transient dynamics [55]. For the inhibitory synaptic weights, we found a regime of parameters yielding functional memory dynamics. This regime seems to correspond to ‘loosely-balanced’ network states, which are characterized by a dynamic equilibrium of excitation and inhibition [20]. In contrast to that, ‘tightly-balanced’ network states feature inhibition that closely follows excitation, and in some cases seem to enable more efficient coding. By introducing inhibitory plasticity [78], our network could possibly enter a tightly-balanced state. This would increase the complexity of our model tremendously, but could be subject to future studies on more efficient memory encoding.

Synaptic plasticity such as long-term potentiation is widely considered to be the main mechanism that underlies memory, as it is expressed by the synaptic plasticity and memory hypothesis [2, 48]. Long-term potentiation of synapses (LTP) has been subdivided into three different types: LTP1, which depends on stimulation only, LTP2, which depends on translation of existent mRNA, and LTP3, which depends on transcription in the soma [2, 3, 62]. LTP2 and LTP3 are often subsumed as late-phase LTP, whereas LTP1 is called early-phase LTP [2, 28], which paradigm is followed by our model. Similar phenomena and processes as the ones discussed for LTP are found for long-term depression (LTD) [2, 69, 75]. In addition to synaptic plasticity, there are hints to other, cell-intrinsic mechanisms that might be relevant for the storage of memory [2], such as non-coding RNAs that produce learning-related epigenetic changes in neurons [40]. However, the potential effects of such non-synaptic memory mechanisms on the results of our study and on memory consolidation in general remain unknown and require further investigations.

Following their encoding, memories can become consolidated to last for hours, days, or even years, depending on conditions like strength of the learning stimulus and neuromodulation [3, 22, 49]. In principal, one can distinguish two different paradigms related to memory consolidation. On the level of brain areas, systems consolidation describes the transfer of newly formed memories, mainly from the hippocampus to the neocortex [22, 49], as it was first discovered in the case of the patient H.M. [71]. This type of consolidation is related to sleep and rest [51, 64]. On the other hand, there is synaptic consolidation, which denotes the in-place stabilization of changes in the strength of synaptic connections [23, 49], and thereby is a synonym for LTP2 and LTP3, or late-phase LTP as discussed before [2]. It is sometimes suggested to use the term cellular consolidation instead of synaptic consolidation [22, 65], since the processes involved are not only located at the synaptic site. This is mostly due to the fact that mRNA and proteins are synthesized in the soma, from where they are transported to the dendrites and spines [3, 17, 22, 65]. Nevertheless, there can also be local protein synthesis in the dendrites [14, 37, 47, 59], which could confine the here identified protein-dependent dynamics of consolidation and improvement to specific dendritic branches of a neuron.

The most important finding of our study is that for many parameter settings, the quality of recall eight hours after learning is much better than the quality of recall 10 seconds after learning. In psychological literature, such an increase in recall quality over time is referred to as memory improvement or hypermnesia [26]. It has been found particularly, but not exclusively, in picture memory [26, 61, 79]. In our simulations and analyses, we found the improvement effect already to occur at the very first recall attempt, while in psychological experiments the effect of improvement has mostly been measured over the course of several recall attempts. Our model accounts as well for improvement by multiple recalls, as we showed using a protocol with an intermediate recall stimulus. Nevertheless, there are experiments indicating that recall can already be improved in the first attempt [26]. It would be desirable to have more experimental data on very early recall as well as on recall after eight hours without previous recall.

Our model explains another interesting, seemingly fundamental feature of memory consolidation. As studies on positive effects of memory disuse have proposed [11, 38], learning will be most powerful after a break, when the storage strength is still quite low and the retrieval strength has again decreased. Attempts to retrieve will then result in a strong increase in storage strength. If the retrieval strength is still high, the gain in storage strength upon this (‘easy’) retrieval will be low. This theory seems to be partially consistent with our results, considering that the storage strength corresponds to the late-phase synaptic weight, and retrieval strength to the early-phase synaptic weight. In our model, increases in early-phase weights upon stimulation are much larger if the early-phase weight is low before the stimulation than if it is high. Since increases in early-phase weight lead to changes in late-phase weight, the gain in late-phase weight (i.e., presumably, the storage strength) indeed inversely depends on the early-phase weight (retrieval strength) before stimulation. Nevertheless, further investigations would have to resolve how our model could match the psychological finding that retrieval at small retrieval strength should take long [11, 38], while at the present our model exhibits fast recall even in the case of low early-phase weights.

Behavioral tagging is a phenomenon that could be the behavioral analog of synaptic tagging, describing the finding that the memory of a weak input perceived around the same time as strong novel stimulation becomes consolidated along with the memory of the strong stimulus [7, 52, 53]. Using their theoretical model of a feed-forward network, Ziegler and colleagues [83] provided a connection between STC mechanisms and behavioral tagging focusing on an inhibitory avoidance paradigm. For this, they considered dopamine modulation to model novelty. By extending our model by such dopamine modulation, more experimental data could be matched to provide further evidence for the relatedness of behavioral tagging and synaptic tagging.

Please note that for several potential extensions of our model discussed above, it would be useful to improve the computational efficiency of the numerical simulation. For that, it is very promising to implement our network model on neuromorphic hardware [18, 44]. Neuromorphic hardware enables the simulation of larger networks and thereby the storage of many cell assemblies. In this way, network capacity and the interference between cell assemblies could be investigated. The neuromorphic approach would not only facilitate scientific investigations but, due to its good performance and energy efficiency [44, 81], also offers great opportunities for the technical application of our model. Implemented on neuromorphic hardware, our model could be used for fast, energy-efficient self-improvement of information storage after a few presentations. In conclusion, our theoretical model provides a further piece of evidence that the STC mechanisms are essential for memory dynamics, and it predicts that these STC mechanisms also allow the improvement of memories, which can be beneficial for biological as well as artificial memory systems.

## Methods

### Model

To simulate the dynamics of memory representations in the hippocampus, we use a network model that comprises spiking neurons and synapses with detailed plasticity features. In the following, we present our mathematical description of neurons and synapses, which is depicted in Fig. 1a. After that, we explain how our network is structured at the population level. The parameters we use are given in Tables 1 and 2.

**Table 1:**
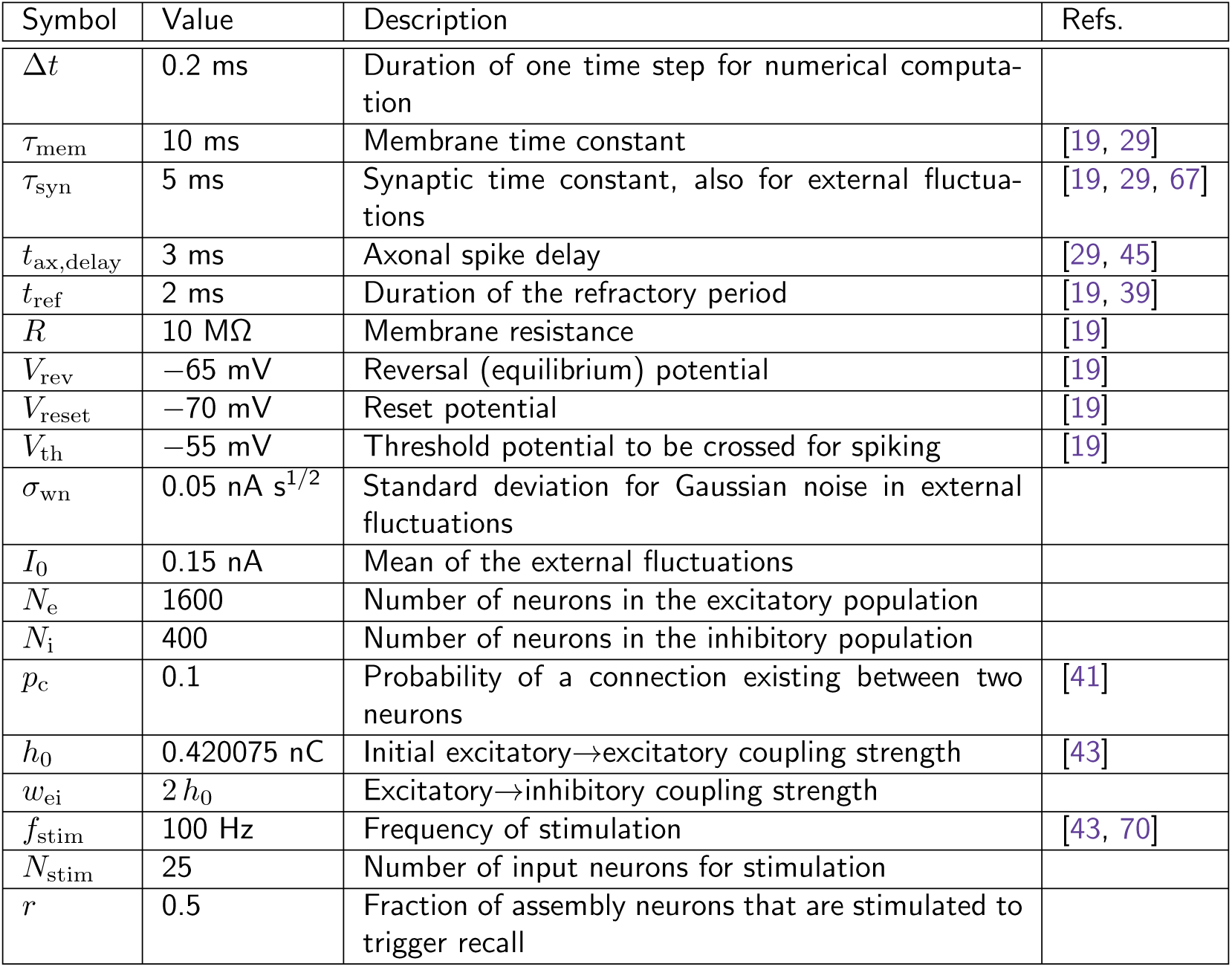
Parameters for neuron and static network dynamics. Values were used as given in this table, unless stated otherwise.

| Symbol | Value | Description | Refs. |
| --- | --- | --- | --- |
| $\Delta t$ | 0.2 ms | Duration of one time step for numerical computation | |
| $\tau_{\text{mem}}$ | 10 ms | Membrane time constant | [19, 29] |
| $\tau_{\text{syn}}$ | 5 ms | Synaptic time constant, also for external fluctuations | [19, 29, 67] |
| $t_{\text{ax,delay}}$ | 3 ms | Axonal spike delay | [29, 45] |
| $t_{\text{ref}}$ | 2 ms | Duration of the refractory period | [19, 39] |
| $R$ | 10 MΩ | Membrane resistance | [19] |
| $V_{\text{rev}}$ | −65 mV | Reversal (equilibrium) potential | [19] |
| $V_{\text{reset}}$ | −70 mV | Reset potential | [19] |
| $V_{\text{th}}$ | −55 mV | Threshold potential to be crossed for spiking | [19] |
| $\sigma_{\text{wn}}$ | 0.05 nA s<sup>1/2</sup> | Standard deviation for Gaussian noise in external fluctuations | |
| $I_0$ | 0.15 nA | Mean of the external fluctuations | |
| $N_{\text{e}}$ | 1600 | Number of neurons in the excitatory population | |
| $N_{\text{i}}$ | 400 | Number of neurons in the inhibitory population | |
| $p_{\text{c}}$ | 0.1 | Probability of a connection existing between two neurons | [41] |
| $h_0$ | 0.420075 nC | Initial excitatory→excitatory coupling strength | [43] |
| $w_{\text{ei}}$ | 2 $h_0$ | Excitatory→inhibitory coupling strength | |
| $f_{\text{stim}}$ | 100 Hz | Frequency of stimulation | [43, 70] |
| $N_{\text{stim}}$ | 25 | Number of input neurons for stimulation | |
| $r$ | 0.5 | Fraction of assembly neurons that are stimulated to trigger recall | |

**Table 2:** Parameters for synaptic plasticity. Values were used as given in this table, unless stated otherwise.

| Symbol | Value | Description | Refs. |
| --- | --- | --- | --- |
| $t_{c,\text{delay}}$ | 0.0188 s | Delay of postsynaptic calcium influx after presynaptic spike | [33] |
| $c_{\text{pre}}$ | 1 | Presynaptic calcium contribution | [33, 43] |
| $c_{\text{post}}$ | 0.2758 | Postsynaptic calcium contribution | [33, 43] |
| $\tau_c$ | 0.0488 s | Calcium time constant | [33, 43] |
| $\tau_h$ | 688.4 s | Early-phase time constant | [33, 43] |
| $\tau_p$ | 60 min | Protein time constant | [17, 43] |
| $\tau_z$ | 60 min | Late-phase time constant | [17, 43] |
| $\gamma_p$ | 1645.6 | Potentiation rate | [33, 43] |
| $\gamma_d$ | 313.1 | Depression rate | [33, 43] |
| $\theta_p$ | 3 | Calcium threshold for potentiation | [43] |
| $\theta_d$ | 1.2 | Calcium threshold for depression | [43] |
| $\sigma_{\text{pl}}$ | 0.290436 nC | Standard deviation for plasticity fluctuations | [33, 43] |
| $\alpha$ | 1 | Protein synthesis rate | [17, 43] |
| $\theta_{\text{pro}}$ | 0.210037 nC | Protein synthesis threshold | [43] |
| $\theta_{\text{tag}}$ | 0.0840149 nC | Tagging threshold | [43] |

### Neuron model

The dynamics of the membrane potential *V*_*i*_(*t*) of the Leaky Integrate-and-Fire neuron *i* is described by [29]:

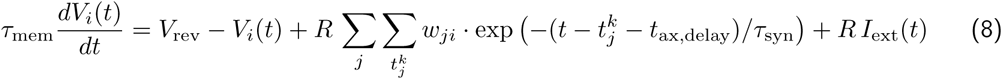

with reversal potential *V*_rev_, membrane time constant *τ*_mem_, membrane resistance *R*, synaptic weights *w*_*ji*_, spike times 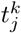, axonal delay time *t*_ax,delay_, synaptic time constant *τ*_syn_, and external input current *I*_ext_(*t*). If *V*_*i*_ crosses the threshold *V*_th_, a spike is generated. The spike time 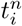 is then stored and the membrane potential is reset to *V*_reset_, where it remains for the refractory period *t*_ref_. The membrane potential dynamics is mainly driven by an external input current that accounts for synaptic inputs from outside the network, described by an Ornstein-Uhlenbeck process:

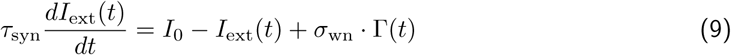

with mean current *I*_0_ and white-noise standard deviation *σ*_wn_. Note that in this equation, Γ(*t*) is Gaussian white noise with mean zero and variance 1*/dt* that approaches infinity for *dt*→ 0. The Ornstein-Uhlenbeck process has the same colored-noise power spectrum as the fluctuating input to cortical neurons coming from a large presynaptic population [21]. Therefore, it is well-suited to model background noise in our model.

### Model of synapses and STC

If there is a synaptic connection from neuron *j* to neuron *i*, all spikes *k* that occur in *j* are transmitted to *i*. The postsynaptic current caused by a presynaptic spike depends on the weight of the synapse. The total weight or strength of a synaptic connection from neuron *j* to neuron *i* is given by:

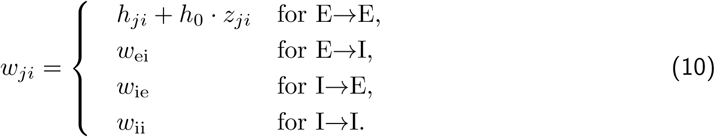

In our model, all synaptic connections involving inhibitory neurons are constant. The weight of E→E connections, however, consists of two variable contributions providing the core of the STC mechanisms: the early-phase weight *h*_*ji*_, and the late-phase weight *z*_*ji*_. The dynamics of the early-phase weight is given by

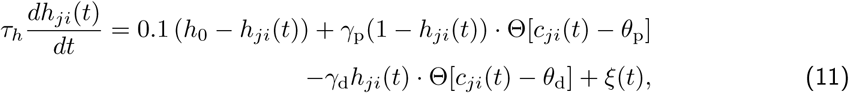

where Θ[·] is the Heaviside function, *τ*_*h*_ is a time constant, and *c*_*ji*_(*t*) is the calcium concentration at the postsynaptic site. The first term on the right-hand side describes a relaxation of the early-phase weight to its initial value *h*_0_, the second term describes early-phase LTP with rate *γ*_p_ for calcium above the threshold *θ*_p_, the third term describes early-phase LTD with rate *γ*_d_ for calcium above the threshold *θ*_d_, and the fourth term 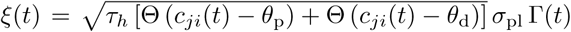 describes calcium-dependent noise-driven fluctuations with standard deviation *σ*_pl_ and Gaussian white noise Γ(*t*) with mean zero and variance 1*/dt*. The calcium concentration *c*_*ji*_(*t*) at the postsynaptic site depends on all past pre- and postsynaptic spikes *n* and *m*, respectively:

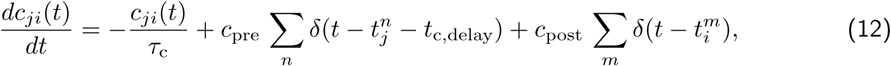

where *δ*(·) is the Dirac delta distribution, *τ*_*c*_ is a time constant, *c*_pre_ is the contribution of presynaptic spikes, *c*_post_ is the contribution of postsynaptic spikes, 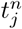 and 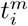 are spike times, and *t*_c,delay_ is the delay of calcium triggered by presynaptic spikes.

The calcium-based plasticity model (Eqs. 11 and 12) that we use to describe the early phase of long-term potentiation and depression, is based on previous theoretical studies [33, 36, 43]. Similar to Li et al. [43], the first term on the right-hand side of Eq. 11 describes a relaxation to the initial condition to which the early-phase weight returns or decays on a timescale of a few hours. This decay accounts for the fact that early-phase changes are transient providing an accurate description of the experimentally verified dynamics of the STC mechanisms [28].

The calcium parameters provided by Graupner and Brunel [33] were obtained by fitting experimental data from in vitro experiments [82]. Since extracellular calcium concentrations are much lower in vivo than in vitro, the parameters need to be corrected for modeling in vivo networks. Following Higgins et al. [36], the calcium influx into the postsynaptic spine can be assumed to decrease proportionally to the ratio of in vivo and in vitro extracellular calcium concentrations, which leads to a factor of 0.6. Therefore, we adjust the values provided by Graupner and Brunel [33] with this factor to use them in our network model.

Driven by the calcium-based early-phase dynamics, the late-phase synaptic weight is given by

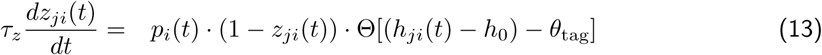

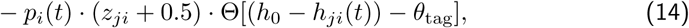

with the protein amount *p*_*i*_(*t*), the late-phase time constant *τ*_*z*_, and the tagging threshold *θ*_tag_. The first term on the right-hand side describes late-phase LTP and the second late-phase LTD. Both depend on the amount of proteins being available. If the early-phase weight change |*h*_*ji*_(*t*) −*h*_0_| exceeds the tagging threshold, the synapse is tagged. This can be the case either for positive or for negative weight changes. The presence of the tag leads to the capture of proteins (if *p*_*i*_(*t*) > 0), and thereby gives rise to changes in the late-phase weight.

The synthesis of new proteins depends on the early-phase weights for increase, but the amount of proteins also inherently decays exponentially:

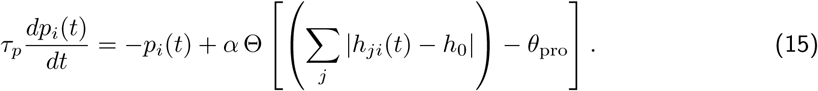

### Population structure

Using the neuron model and the synapse model explained above, we set up a neural network consisting of 1600 excitatory and 400 inhibitory neurons (depicted in Fig. 1b). This ratio between excitatory and inhibitory neurons is commonly used for cortical and hippocampal networks [13]. Some of the excitatory neurons receive specific inputs to learn and recall a memory representation (see next subsection). As described before, only the synapses between excitatory neurons are plastic. The inhibitory population serves to provide realistic feedback inhibition. The overall connectivity across both populations is 10%, meaning that there is a probability of 0.1 that the link from any neuron to another one in the whole network exists. The value is reasonable for hippocampal region CA3 [41].

### Learning and recall procedure

Before we stimulate our network, we first let the initial activity settle for 10.0 seconds. After that, we apply our learning protocol, which delivers three stimulus pulses, of 0.1 seconds duration each, to the neurons belonging to the desired cell assembly (for instance, to the first 150 neurons in the network). The pulses in our protocol are separated by breaks of 0.4 seconds. The stimulation is delivered by an Ornstein-Uhlenbeck process (see Eq. 9 above) whose mean is the number of putative input neurons *N*_stim_ times the input frequency *f*_stim_. For the reproduction of previous single-synapse investigations ([43], Fig. 2), we used Poisson spikes that were generated using the same number of input neurons and the same stimulation frequency.

After 20.0 seconds, we save the state of the whole network and then apply a recall stimulus with the same input strength as the learning stimulus for 0.1 seconds to half of the neurons in the cell assembly (regarding the example above, to 75 randomly-drawn neurons). We refer to this as “10s-recall”. Next, we load the previously saved network state, such that the network is back in the state it was in immediately before recall. This time, we let the network run for 28810.0 seconds until we apply a recall stimulus, which we refer to as “8h-recall”. For one part of our study, we apply another, intermediate, recall stimulus at a time in between 10 s and 8 h. Following this intermediate recall, we do not reset the network by loading an earlier state, such that it in fact affects the later 8h-recall.

### Measures of recall performance

To investigate the effects of recall stimulation, we divide the excitatory population into thre subpopulations: control neurons that are not directly stimulated neither by learning nor by recall stimulation, cell assembly neurons that are stimulated by both learning and recall stimulus, and cell assembly neurons that are stimulated during learning but not during recall (cf. Fig. 3d). The mean activities of these three subpopulations are given by:

- 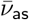: mean activity of the neurons stimulated by the learning stimulus and the recall stimulus (as = “assembly, stimulated”),
- 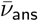: mean activity of the neurons stimulated by the learning stimulus but not by the recall stimulus (ans = “assembly, not stimulated”),
- 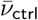: mean activity of the neurons stimulated by neither of the two stimuli (ctrl = “control”).

Based on these mean activities, the recall quality can be measured by computing a quality coefficient:

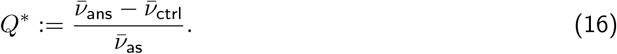

The coefficient typically lies within the interval (0, 1). To achieve pattern completion, the non-stimulated assembly neurons have to be indirectly activated following the activation of the rest of the core neurons, while control neurons should remain relatively silent:

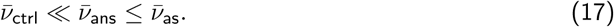

Hence, the value of *Q*^*\**^ will be significantly larger than zero or even approach unity for good pattern completion, that is, for good recall. On the other hand, if it approaches zero, there is either no pattern completion, which means that 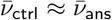, or the network activity diverges, which means that 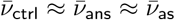.

Even if for one trial *Q*^*\**^ ≫ 0, the pattern completion effect is not necessarily robust across trials for every parameter setting. Therefore, to ensure significant pattern completion, we average over trials:

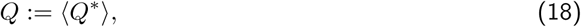

For Fig. 4, due to the lack of error bars, we have to indicate the cases in which no robust pattern completion occurs. Thus, we use the following conditional definition for the quality coefficient:

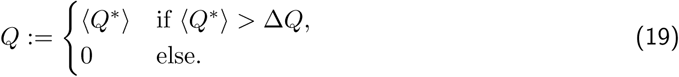

Put in words, the robustness criterion requires the mean ⟨*Q*^*\**^⟩ to be non-negative, and its absolute value to be larger than the standard deviation Δ*Q*. If this is not fulfilled, pattern completion is assumed absent and *Q* is set to zero.

In addition to the subpopulation-based quantity *Q*, we measure the mutual information *MI*_*v*_ between the activity distribution during the recall phase and the activity distribution during the learning phase. The mutual information does not directly relate to pattern completion, but it has the advantage that it is independent of any predefined patterns such as the learning stimulus.

The mutual information of the activity distribution is calculated from the entropy during the learning phase at time *t*_learn_ = 11.0 s, the entropy during the recall phase at time *t*_recall_ ∈ {20.1 s, 28810.1 s}, and the joint entropy between both:

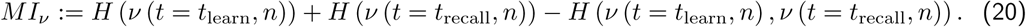

The firing rate function *v* (*t, n*) returns the firing rate of a given neuron *n* at a given time *t*, computed using a sliding window of 0.5 s. To achieve better statistics for our results, we also average the mutual information value over multiple trials and use the standard deviation as error.

### Computational implementation and software used

We used `C++` in the ISO 2011 standard to implement our simulations. To compile and link the code, we employed `g++` in version 7.4.0 with `boost` in version 1.65.1. Our code will be released under an open-source license upon publication of the article.

Random numbers were generated using the generator `minstd_rand0` from the C`++` standard library, while the system time served as the seed. We implemented a loop in our code which ensured that for each distribution a unique seed was used. Unless stated otherwise, we ran each simulation ten times to obtain mean and error estimates of the respective quantities.

For the creation of plots we used `gnuplot` 5.0.3, as well as `Matplotlib` 2.0.0 with `Python` 3.6.8 and `NumPy` 1.16.4.

The network simulations that we needed to perform for this study were computationally extremely demanding. Fortunately, we had the opportunity to use the computing cluster of the *Gesellschaft für wissenschaftliche Datenverarbeitung mbH Göttingen (GWDG)*, which enables fast computation on a large set of processing units. However, despite this strong computational power and the usage of compiled C`++` code, running our spiking network simulations in full detail still took unacceptably long. Thus, to be able to simulate our network faster, we used an approximation that neglects the spiking dynamics in periods without external stimulation. In these periods, we just computed the late-phase dynamics and the exponential decay of the early-phase weights. Supplementary Fig. S2 shows that this approach is justified because the weight dynamics of a synapse does not change if sparsely occurring spikes are neglected. Furthermore, the mutual information conveyed by 8h-recall is in the same regime for full and approximating computation.

## Acknowledgments

We would like to thank the members of the Department of Computational Neuroscience for many fruitful discussions and various helpful comments on this study. The research was funded by the German Research Foundation (CRC1286, project C1, project #419866478) and by the H2020 – FETPROACT project Plan4Act (#732266).

## Author contributions

Jannik Luboeinski: conceptualization; data curation; formal analysis; investigation; methodology; software; validation; visualization; writing and editing.

Christian Tetzlaff: conceptualization; funding acquisition; investigation; methodology; project administration; resources; supervision; writing and editing.

