## Supplementary Information for "Memory consolidation and improvement by synaptic tagging and capture in recurrent neural networks"


8 May 2020

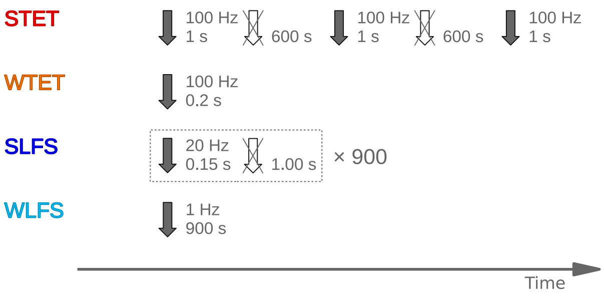


Figure S1: Standard protocols for the the induction of early- and late-phase synaptic potentiation and depression as used in [1].

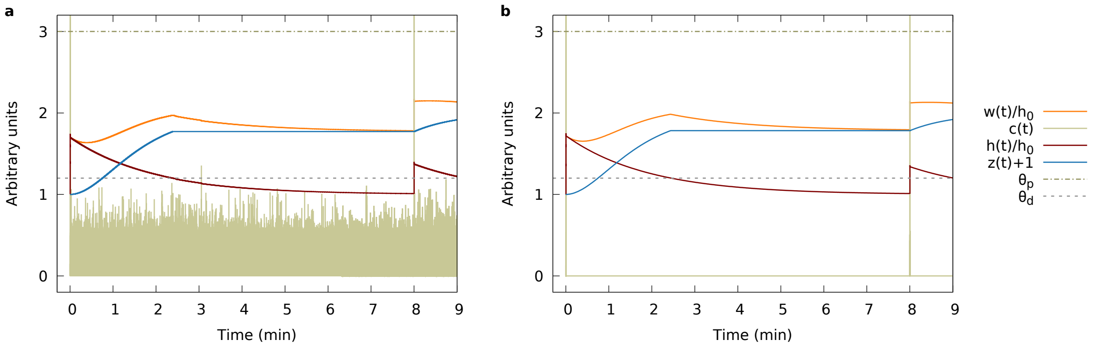


Figure S2: Matching temporal dynamics of the weight of a synapse in **(a)** a spiking network simulation run in full detail and **(b)** an accelerated network simulation with spiking dynamics only during relevant learning- and retrieval-related periods. Outside of these periods, the fully-simulated calcium concentration stays below the plasticity thresholds and thus, can be neglected. Parameter setting: $w_{\mathrm{ie}}/h_0 = 4$, $w_{\mathrm{ii}}/h_0 = 4$, $n_{\mathrm{CA}} = 150$.

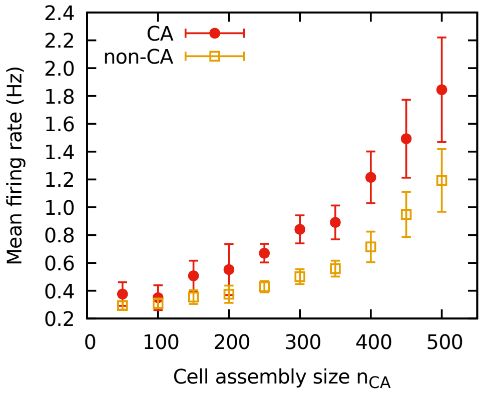


Figure S3: The mean standby firing rate in the excitatory population of a network holding one cell assembly grows strictly monotonically with the size of the assembly (measurement 10 seconds after learning the assembly). Firing rates within the assembly (CA) are higher than in the rest of the population (non-CA). The firing rates for small- to medium-sized assemblies lie in the typical physiological range for the hippocampus of $0.5–1.0\ \mathrm{Hz}$ [2]. Error bars indicate the standard deviation across the respective subpopulation. Parameter setting: $w_{\mathrm{ie}}/h_0 = 4$, $w_{\mathrm{ii}}/h_0 = 4$.

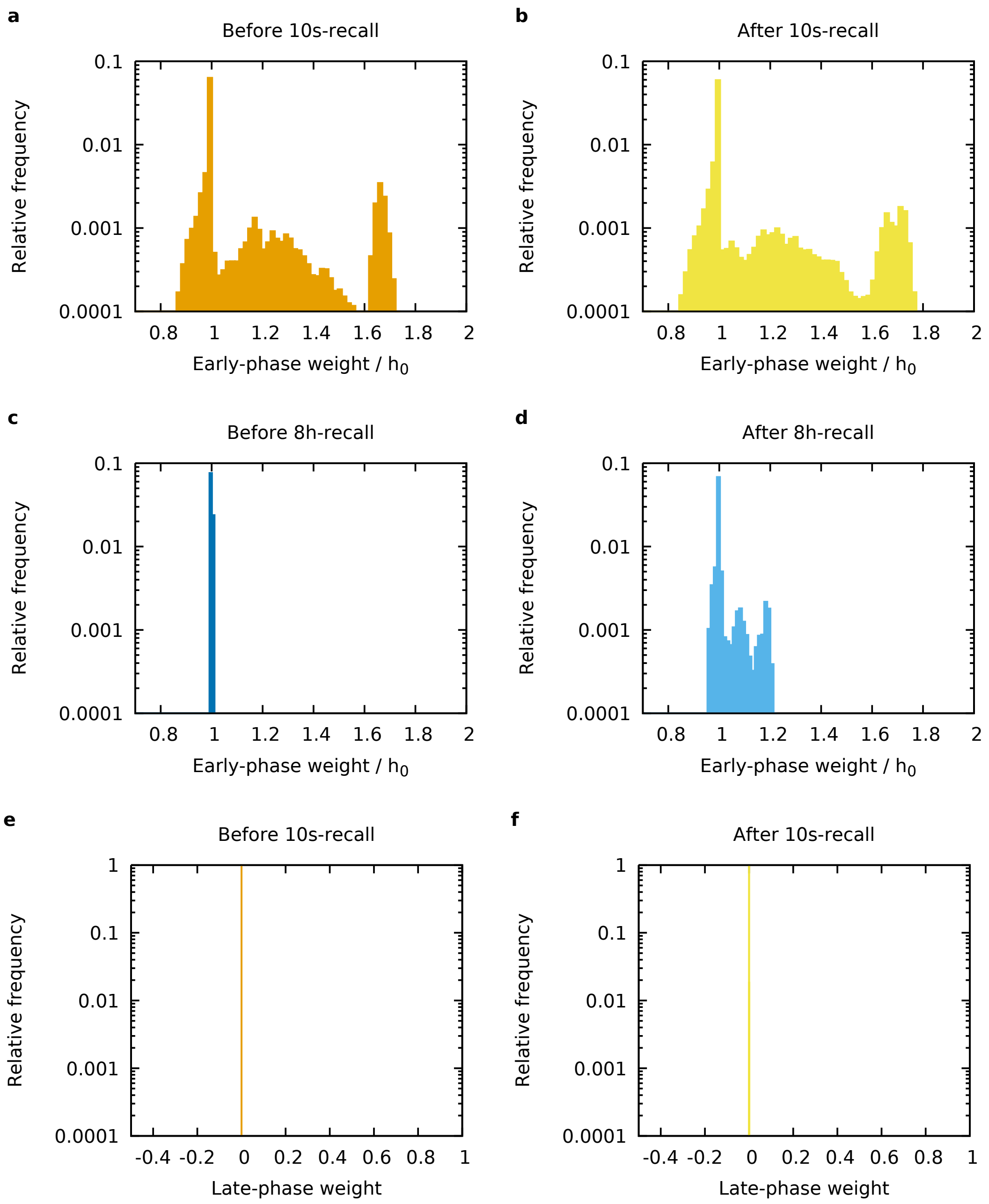
a
Before 10s-recall
Relative frequency
0.1
0.01
0.001
0.0001
0.8 1 1.2 1.4 1.6 1.8 2
Early-phase weight / $h_0$
b
After 10s-recall
Relative frequency
0.1
0.01
0.001
0.0001
0.8 1 1.2 1.4 1.6 1.8 2
Early-phase weight / $h_0$
c
Before 8h-recall
Relative frequency
0.1
0.01
0.001
0.0001
0.8 1 1.2 1.4 1.6 1.8 2
Early-phase weight / $h_0$
d
After 8h-recall
Relative frequency
0.1
0.01
0.001
0.0001
0.8 1 1.2 1.4 1.6 1.8 2
Early-phase weight / $h_0$
e
Before 10s-recall
Relative frequency
1
0.1
0.01
0.001
0.0001
-0.4 -0.2 0 0.2 0.4 0.6 0.8 1
Late-phase weight
f
After 10s-recall
Relative frequency
1
0.1
0.01
0.001
0.0001
-0.4 -0.2 0 0.2 0.4 0.6 0.8 1
Late-phase weight

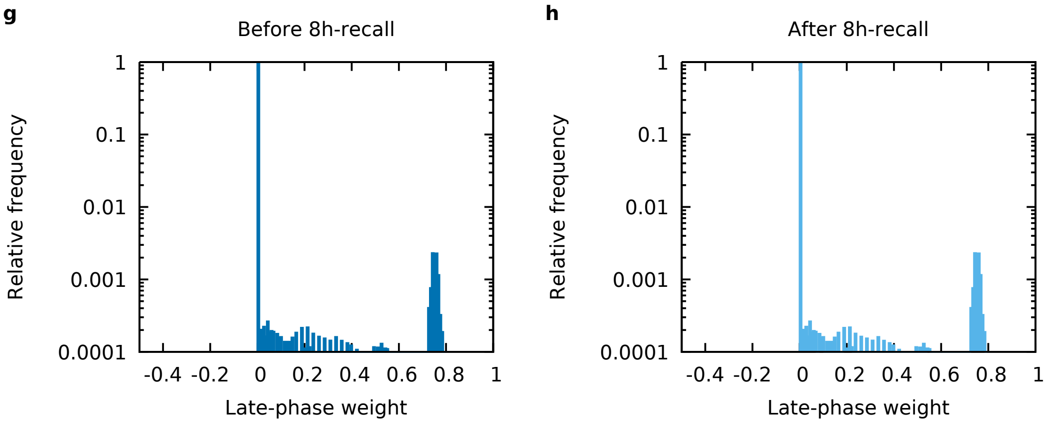


Figure S4: 10s-recall leaves the early-phase weights almost unchanged **(a,b)**, while 8h-recall causes a significant change in the distribution of early-phase weights **(c,d)**. Due to their long timescales, the late-phase weights are not directly affected by recall. There is no substantial late-phase change after $10$ seconds **(e,f)**, but after $8$ hours, late-phase weights are significantly elevated **(g,h)**. The weights in each plot were discretized into $100$ bins. Note that the normalized initial value of early-phase weights is unity and the initial value of late-phase weights is zero.
